# Complete functional mapping of infection- and vaccine-elicited antibodies against the fusion peptide of HIV

**DOI:** 10.1101/307587

**Authors:** Adam S. Dingens, Priyamvada Acharya, Hugh K. Haddox, Reda Rawi, Kai Xu, Gwo-Yu Chuang, Hui Wei, John R. Mascola, Bridget Carragher, Clinton S. Potter, Julie Overbaugh, Peter D. Kwong, Jesse D. Bloom

## Abstract

Eliciting broadly neutralizing antibodies (bnAbs) targeting envelope (Env) is a major goal of HIV vaccine development, but cross-clade breadth from immunization has only sporadically been observed. Recently, Xu et al (2018) elicited cross-reactive neutralizing antibody responses in a variety of animal models using immunogens based on the epitope of bnAb VRC34.01. The VRC34.01 antibody, which was elicited by natural human infection, targets the N terminus of the Env fusion peptide, a critical component of the virus entry machinery. Here we precisely characterize the functional epitopes of VRC34.01 and two vaccine-elicited murine antibodies by mapping all single amino-acid mutations to the BG505 Env that affect viral neutralization. While escape from VRC34.01 occurred via mutations in both fusion peptide and distal interacting sites of the Env trimer, escape from the vaccine-elicited antibodies was mediated predominantly by mutations in the fusion peptide. Cryo-electron microscopy of four vaccine-elicited antibodies in complex with Env trimer revealed focused recognition of the fusion peptide and provided a structural basis for development of neutralization breadth. Together, these functional and structural data suggest that the breadth of vaccine-elicited antibodies targeting the fusion peptide can be enhanced by specific interactions with additional portions of Env. Thus, our complete maps of viral escape provide a template to improve the breadth or potency of future vaccine-induced antibodies against Env’s fusion peptide.

**Author summary:** A major goal of HIV-1 vaccine design is to elicit antibodies that neutralize diverse strains of HIV-1. Recently, some of us elicited such antibodies in animal models using immunogens based on the epitope of a broad antibody (VRC34.01) isolated from an infected individual. Further improving these vaccine-elicited antibody responses will require a detailed understanding of how the resulting antibodies target HIV’s envelope protein (Env). Here, we used mutational antigenic profiling to precisely map the epitope of two vaccine-elicited antibodies and the template VRC34.01 antibody. We did this by quantifying the effect of all possible amino acid mutations to Env on antibody neutralization. Although all antibodies target a similar region of Env, we found clear differences in the functional interaction of Env with the vaccine- and infection-elicited antibodies. We combined these functional data with structural analyses to identify antibody–Env interactions that could improve the breadth of vaccine-elicited antibodies, and thereby help to refine vaccination schemes to achieve broader responses.

## Introduction

The isolation of broadly neutralizing antibodies (bnAbs) capable of neutralizing diverse clades of HIV-1 has invigorated hopes of developing a broadly protective antibody-based HIV vaccine [1,2]. Epitope mapping of bnAbs has revealed a number of conserved sites of vulnerability on Env, and designing immunogens to elicit antibodies that target such sites is a promising vaccine strategy [3,4]. Structural characterization of the epitope of bnAb N123-VRC34.01 (subsequently referred to as VRC34.01) revealed that the N terminus of the fusion peptide (FP) is one such site of vulnerability [5]. A number of additional bnAbs that also partially target this epitope have been characterized [6–10], suggesting it may be a promising vaccine target.

Xu et al (2018) [11] used the VRC34.01 epitope as a template to design vaccine approaches that elicited antibodies to the fusion peptide capable of neutralizing diverse strains of HIV-1. The broadest of these vaccine-elicited fusion peptide-directed antibodies (vFP antibodies) derive from a class of murine antibodies, the vFP1 class, whose members share similar B cell ontogenies and structural modes of recognition. Here we focus primarily on two of these antibodies, 2717-vFP16.02 and 2716-vFP20.01, which have breadths of 31% and 27% respectively [11]. The vFP16.02 antibody was elicited by a BG505 Env trimer prime and a FP-carrier protein conjugate (FP-KLH) boost, and vFP20.01 was elicited via a similar scheme with an additional timer boost. The breadth of these antibodies rivals infection-elicited bnAbs such as 2G12 [12], although the vFP antibodies still have lower affinity for Env than the original VRC34.01 bnAb [11]. To improve breadth and potency in further rounds of structure-based vaccine design, we undertook studies to precisely define the epitope specificities of the vFP antibodies and to understand how they differ from the original template VRC34.01 bnAb.

## Results

We used mutational antigenic profiling [13] to quantify how all single amino-acid mutations at each residue of Env affected the neutralization of replication-competent HIV by VRC34.01 and the two vaccine-induced antibodies, vFP16.02 and vFP20.01. Specifically, for each antibody, we neutralized libraries [14] of viruses carrying all amino-acid mutations to the ectodomain and transmembrane domain of the BG505.T332N Env at an ~IC_95_ antibody concentration. For each of the possible 12,730 Env mutations (670 mutagenized sites × 19 amino acid mutations), we calculated the enrichment of the mutation in the antibody-selected condition relative to a non-selected mutant virus library, a quantity that we term *differential selection* [13] (Fig S1A). This entire process was performed in full biological replicate, beginning with independent generation of the proviral plasmid mutant libraries. The results from the two replicates were well correlated (Fig S1B, S1C). For the remainder of the paper, we present the averaged data from these two replicates.

It was immediately apparent that viruses with mutations in the N terminus of the fusion peptide (sites 512-516) robustly escaped all three antibodies (Fig 1A, S2–4). However, there were also obvious differences between VRC34.01 and the two vaccine-elicited antibodies. While viral escape was concentrated in the fusion peptide for the vFP antibodies, there were a number of additional sites in Env where mutations allowed for escape from VRC34.01. Prior structural and functional characterization of VRC34.01 have revealed that it interacts with both the N terminus of the fusion peptide and the N-linked glycan at residue 88 [5]. The results of these prior structural and functional studies are highly correlated with our mutational antigenic profiling data (Fig S2, S5). Mutational antigenic profiling also revealed additional sites of viral escape from VRC34.01, including residues 84, 85, 227, 229, 241 and 243, which are all proximal to the fusion peptide (Fig 1B). Interestingly, we also observed a modest but reproducible signal for escape from vFP16.02 and vFP20.01 via introducing a glycosylation motif at N241, where a glycan would cause steric hindrance to the approach of these antibodies, and via mutations at site 85 (Fig 2, S3, S4).

**Fig 1.**
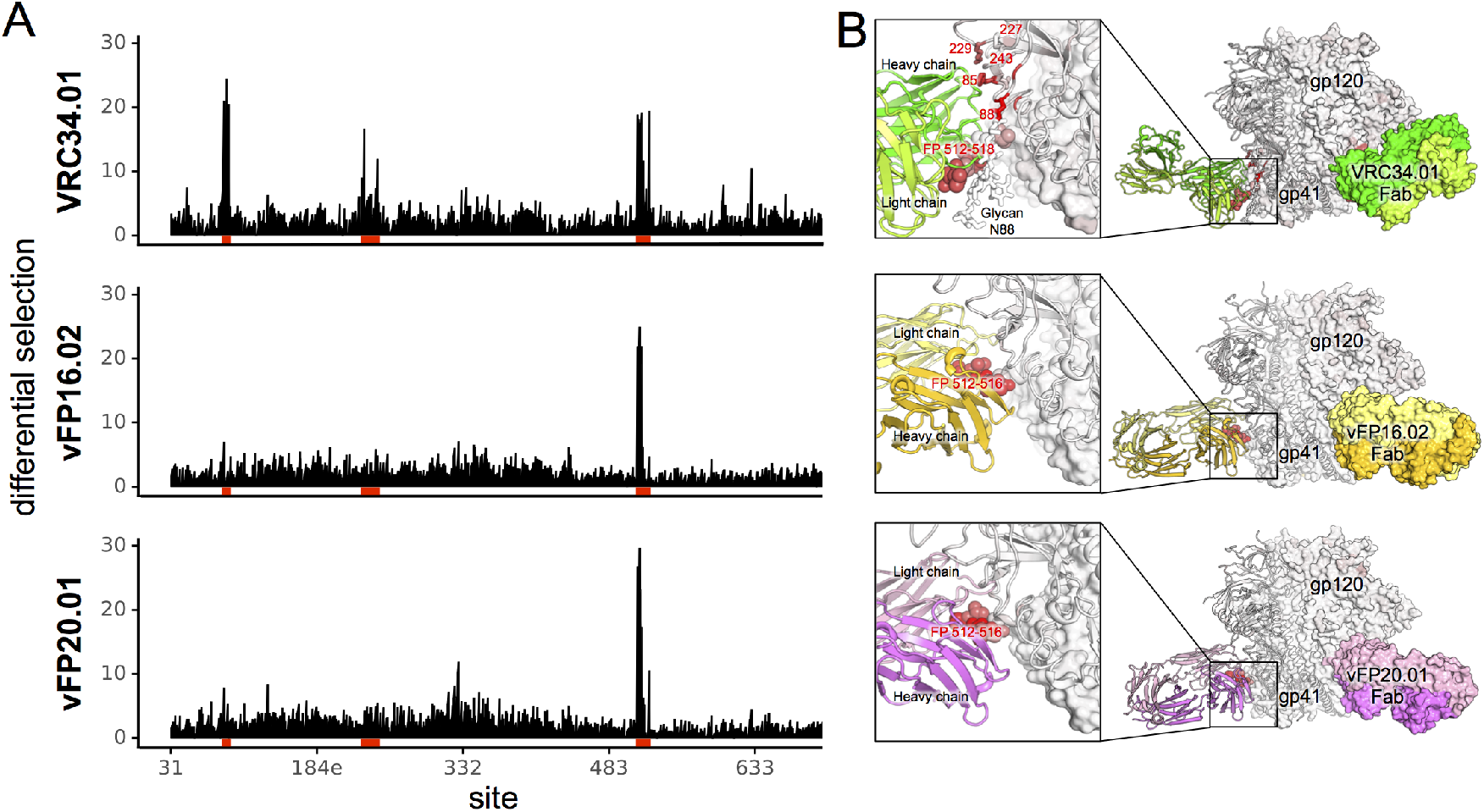
Complete escape profile of natural and vaccine elicited fusion peptide antibodies. **A.** The positive site differential selection is plotted across the length of the mutagenized portion of Env for each antibody. **B.** Structural representation of the FP-directed antibody epitope on BG505 Env trimer. The Env trimer is colored from white to red according to the positive differential selection at each site. Cartoon diagram of Fab-trimer complexes were adapted from crystal structure (VRC34.01) and cryoEM structure (vFP16.02 and vFP 20.01), with one Fab-protomer shown in ribbon and the other two Fab-protomers shown as surface. Labeled fusion peptide residues are shown in spheres. Detail of the Fab-trimer interaction was shown in the inset.

**Fig 2.**
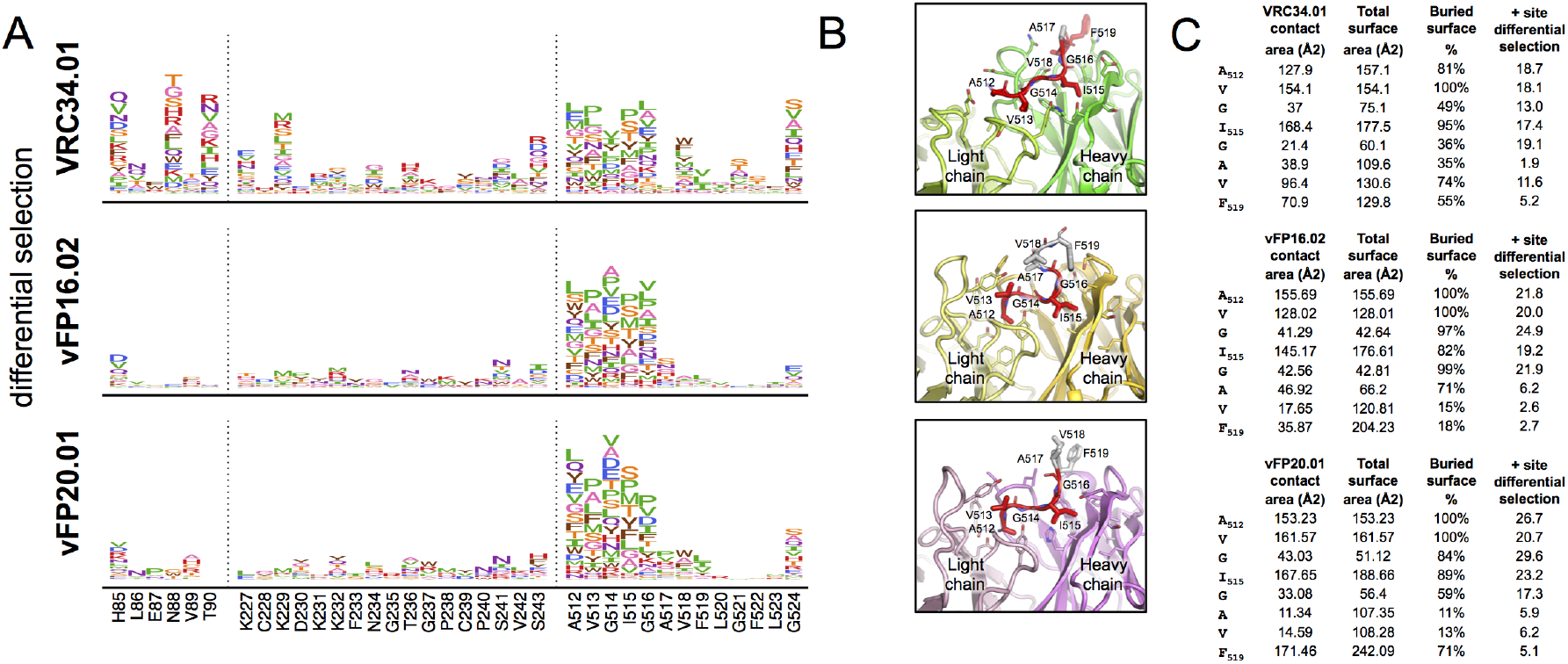
The mutation-level escape profiles agree with structural characterizations of the fusion peptide epitope. **A.** The mutation-level escape profile for regions with considerable differential selection by at least one antibody (regions with red underlay in Fig 1A). Left to right, these include the N88 glycosylation motif and neighboring area, surface exposed gp120 residues that interact with VRC34.01, and most of the fusion peptide. The height of each amino acid is proportional to the logarithm of the relative enrichment of that mutation in the antibody selected condition relative to the non-selected control. **B.** Fab-FP peptide crystal structures of each antibody. Antibody Fabs were colored as in Fig. 1B, FP residues with significant differential selection to each antibody were colored in red. **C.** Total surface, antibody-contact surface (determined from co-crystal structures shown in Fig 2B), and positive site differential selection for each antibody

Since we measured the effect of every amino acid at every site, we can examine viral escape at a *mutation* rather than *site* level (Fig 2A). For example, V518W, V518M, and V518L resulted in drastic escape from VRC34.01 neutralization, while V518A had no effect (Fig S2, S5, and Kong et al (2016) [5]). Further, while the mutational antigenic profiling results agreed with alanine and glycine scans of the fusion peptide [11], the complete escape profiles identified additional escape mutants at nearly all of the fusion peptide residues that contact each antibody. This mutation-level mapping was especially informative for understanding the mechanistic basis for escape from neutralization. For example, hydrophobic amino acids were among the most enriched escape mutants at site 512-516 for all three antibodies. This may reflect their ability to eliminate antibody binding while still allowing for the fusion peptide to be efficiently inserted into the host cell plasma membrane during fusion.

Even within the fusion peptide region, there are differences between the template, infection-elicited bnAb and the vaccine-elicited antibodies. Escape from VRC34.01 could also occur via mutations at sites 518 and 521, whereas these mutations had little impact on the vaccine-elicited antibodies. These residue level differences in escape within the FP agree with structural interactions; within FP peptide-Fab co-crystal structures, the percent buried surface area of each residue tracks well with the extent of escape for each antibody (Fig 2B, C). Likewise, the fusion peptide dominated the epitope of antibodies vFP16.02 and vFP20.01 bound to Env, with the cryo-EM reconstructions showing strongest electron density of the bound antibody around the fusion peptide [11], with residues 512-519 accounting for ~63% and ~56% of the total protein epitope surface for vFP16.02 and vFP20.01, respectively (S1 Tables). However, these data also highlighted how not all structural contacts contribute equally to binding. Cryo-EM (vFP16.02 and vFP20.01) and crystal (VRC34.01) structures of each Fab bound to HIV-1 Env showed contacts of the antibodies with residues 517, 519, and 520 of the fusion peptide of the trimeric Env (Fig 1B, Tables S1 B, C). Mutational antigenic profiling suggested that these residues were less critical to escape than residues 512-516.

We also found an unexpected site of viral escape at site 524 that was shared between all three antibodies. Although Gly 524 is distal to the epitope and does not directly contact any of the antibodies, it is part of a structural motif conserved in all three antibodies, involving a tight turn that allows Phe 522 to anchor the flexible fusion peptide via van der Waals interactions to α6 helix of gp41, and β5 and β7 strands of gp120 (Fig S6) [5,11]. Interestingly, mutations to this site mediated escape from each antibody to differing extents (Fig 1). Thus, these distal mutations likely altered the presentation or conformational dynamics of the N terminus of the fusion peptide, an observation relevant to immunogen design.

Sequence analysis of sensitive and resistant pseudoviruses provides an alternative approach to identifying naturally occurring individual Env mutations that were associated with resistance. Analysis of the FP sequences of a panel of 208 sensitive and resistant pseudoviruses identified mutations at sites 513 and 515 that were associated with resistance to the vFP antibodies, while mutations at sites 513, 514, 518, and 519 were associated with resistance to VRC34.01 (Fig S7). More precisely, in the case of VRC34.01, the sequence-neutralization association analysis revealed that methionine and leucine at position 518 and isoleucine at position 519 were significantly associated with resistance, which were among the escape mutations identified by the antigenic profiling analysis (Fig 2). However, the known critical glycosylation motifs such as N88 or N611 were not detected to be associated with antibody sensitivity in analysis of the pseudovirus panel. This is likely due to the fact that N88 and N611 are highly conserved in the panel of 208 pseudoviruses. While the resistance analysis was limited to mutations present at sufficient numbers in the pseudovirus panel, it nevertheless allowed for an examination of the effect of each mutation in diverse strains. These results agreed with the comprehensive escape profiles and highlighted naturally occurring diversity that may present a challenge to fusion peptide-targeting vaccines.

Mutational antigenic profiling also identified mutations distal to the epitope that were depleted, rather than enriched, during antibody selection relative to the non-selected libraries (Fig 3). These include mutations that eliminated the N611 glycosylation motif, indicating viruses that lack the N611 glycan are more sensitive to neutralization. To a lesser extent, we observed the same phenotype at the N88 glycosylation motif for the vFP antibodies but not for VRC34.01 (Fig 3A, 3B). These results are consistent with the fact that binding of VRC34.01 is dependent on the N88 glycan [5], and disrupting the N88 and N611 glycosylation motifs individually in BG505 pseudoviruses results in more potent neutralization by vFP16.02 and vFP20.01 [11]. Numerous mutations throughout Env were depleted during VRC34.01 selection, while far fewer mutations were depleted during vFP16.02 and vFP20.01 selection (Fig 3A). For all three antibodies, sites 621 and 624 were amongst those under the strongest negative differential selection (Fig 3A, 3B). These surface-exposed gp41 sites, which neighbor the FP epitope but do not directly interact with any of the antibodies, could allosterically affect the conformation and/or presentation of the fusion peptide or neighboring glycans. Indeed, site 621 is part of a surface salt bridge network that could be important for maintaining local structure (Fig S8) [11]. While the mechanism of this phenomenon is yet to be fully elucidated, introducing these mutations into Env trimers may be a way to increase the exposure of the fusion peptide in trimer immunogens.

**Fig 3.**
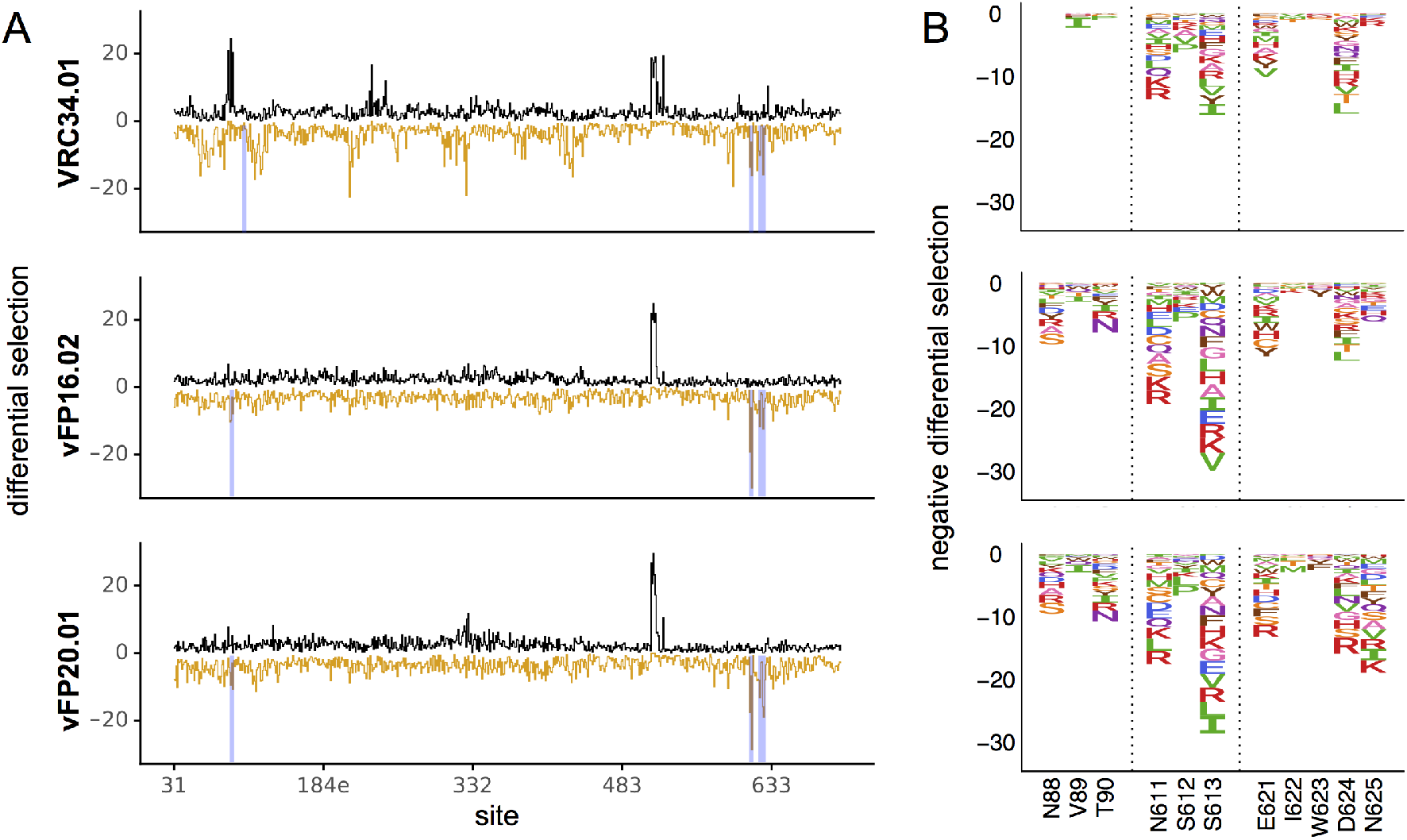
Mutations that better expose the fusion peptide are selected against during antibody treatment. **A.** The positive (black) and negative (orange) site differential selection is plotted across the length of the mutagenized portion of Env for each antibody. **B.** The negative differential selection for regions of interest (regions highlighted in blue in 1A). Left to right, these include the N88 glycosylation motif, the N611 glycosylation, and surface exposed gp41 sites that consistently have mutations depleted upon antibody selection. The height of each amino acid is proportional to the logarithm of the relative depletion of that mutation in the antibody selected condition relative to the non-selected control.

Our mutational antigenic profiling may also help to contextualize prior structural analyses [5,11]. vFP antibodies have a restricted angle of approach and substantial interactions with glycans at residues N88, N295, N448, and N611 [11]. We showed that eliminating these glycans did not result in viral escape, thus indicating that that these interactions do not contribute to binding energetics. Rather, these antibodies likely approach the fusion peptide at an angle that accommodates or avoids steric clashes with glycans.

Further, vFP16.02 and vFP20.01’s focused recognition of the fusion peptide’s N-terminus is also reflected by their Env-bound cryo-EM reconstructions [11]. Reduced electron density for the antibody outside of this region around the fusion peptide indicated greater conformational variability as a result of being less constrained by interactions with Env.

To further understand the structural basis for development of antibody breadth for the vaccine-elicited antibodies, we determined cryo-EM structures of two additional vFP antibodies of the vFP1 class, vFP1.01 and vFP7.04 (8% and 10% breadth, respectively), in complex with BG505 DS-SOSIP Env trimer (Fig 4, Fig S9, Tables S1, Table S2). Using a similar strategy as was adopted for the cryo-EM reconstructions of Env-bound complexes of vFP16.02 and vFP20.01 [11], we obtained cryo-EM reconstructions, at overall resolutions of 4.2 A and 4.0 Å, respectively, of vFP1.01-BG505 DS-SOSIP-VRC03-PGT122 and vFP7.04-BG505 DS-SOSIP-VRC03-PGT122 complexes, where the antigen binding fragments (Fabs) of antibodies PGT122 and VRC03 were added to increase the size of the complex and to provide fiducial markers allowing better particle visualization and alignment.

**Fig 4.**
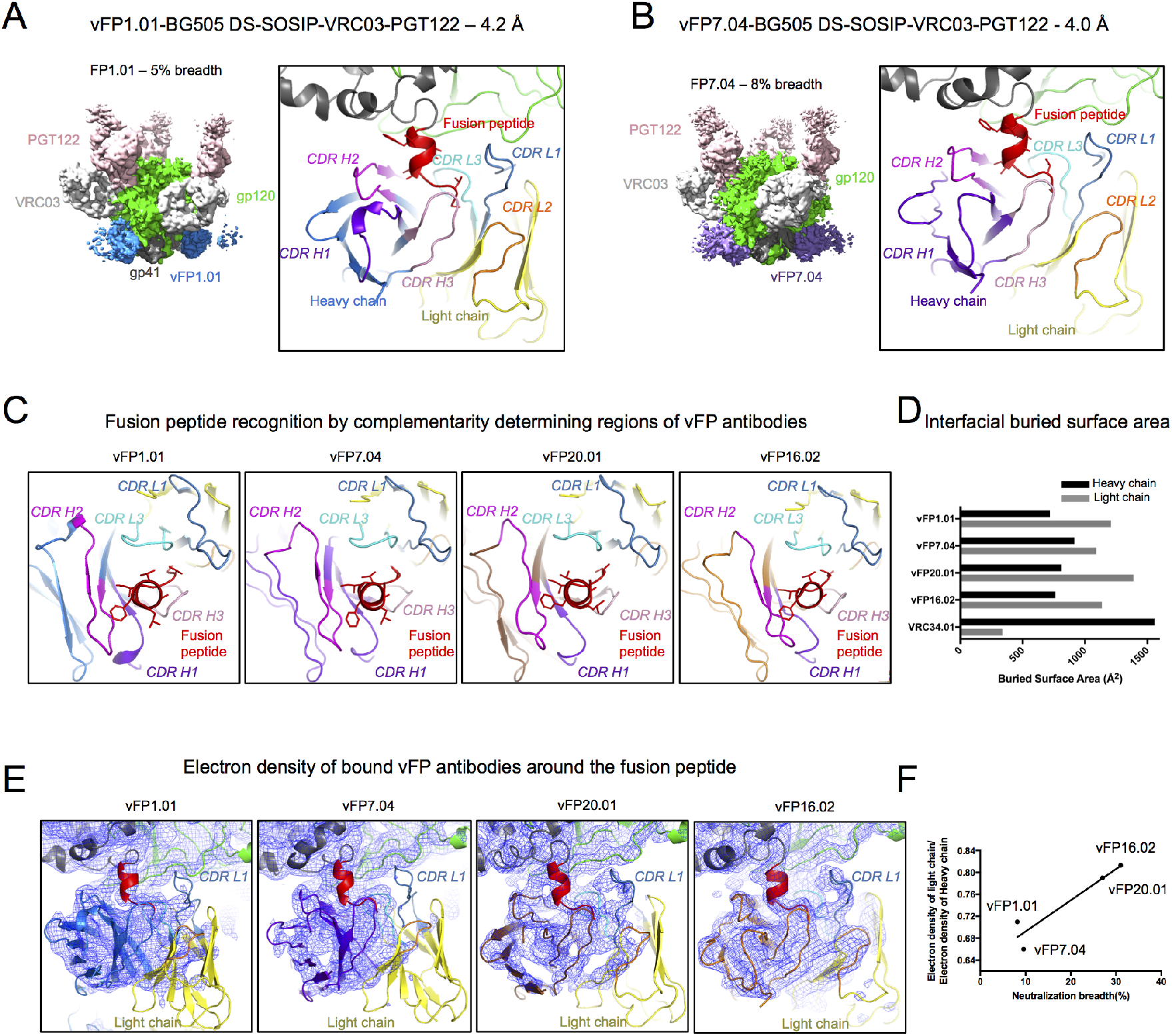
Structural basis for development of neutralization breadth of vFP antibodies. **A.** Left. CryoEM map of quaternary complex with antibody vFP1.01, segmented by components at a contour level that allowed visualization of Fv domains of antibodies. Right. Zoom-in of the fusion peptide region showing vFP1.01 bound with its CDR loops color-coded. **B.** Same as A, but with vFP7.04. **C.** Fusion peptide recognition by vFP1.01, vFP7.04, vFP16.02 and vFP20.01. **D.** Surface area buried at Env interface by the heavy and light chains of vFP1.01, vFP7.04, vFP20.01, vFP16.02 and VRC34. **E.** vFP antibody-bound fusion peptide with electron density from the cryo-EM reconstructions shown as blue mesh. All maps were low pass filtered to 4.5 Å. **F.** Light chain/heavy chain electron density ratio vs. neutralization breadth.

The cryo-EM reconstructions revealed vFP1.01 and vFP7.04 bound to a helical fusion peptide, in what appears to be a signature for this class of vaccine-elicited, murine antibodies (Fig 4, Fig S9, S10). By contrast, VRC34.01 binds an extended conformation of the fusion peptide; this ability to adopt different conformations demonstrates substantial structural plasticity with residue Phe 522 anchoring a highly flexible fusion peptide N terminus to the rest of the Env. In all four vaccine-elicited antibodies, vFP1.01, vFP7.04, vFP16.02 and vFP20.01, a conserved mode of recognition was observed with similar heavy and light chain orientations, with both the heavy and light chains sequestering the fusion peptide N terminus, and the light chain making additional contacts with gp120. Buried surface area calculations using the coordinates fitted and refined into the cryo-EM reconstructions revealed a 1.2-1.7-fold larger interfacial buried surface for the light chain than the heavy chain (Fig 4D). VRC34.01, on the other hand, showed a distinctly different mode of recognition with the heavy chain dominating recognition of the gp41 fusion peptide and gp120, burying ~4.5 fold more interfacial surface area than the light chain (Fig 4D), with the light chain interactions restricted to the N terminal tip of the fusion peptide and glycan 88 that forms a well-ordered interface involving both the heavy and light chains (Table S1 A).

Although the light chain made a larger footprint on Env than the heavy chain in the vFP1.01, vFP7.04, vFP16.02 and vFP20.01 complexes, closer examination of the electron density revealed greater disorder in the light chain density in all the structures. This suggests local variability in the position of the bound light chain that was less constrained by its interactions with Env, whereas the heavy chain made a more stable interface, albeit with a smaller footprint, interacting primarily with the fusion peptide with some interactions with glycan 88. The most stable interface made by the light chain in all four complexes was via the CDR L3 loop interacting with the β0 region of gp120, and contacting gp120 residue His 85 (Fig S11). We observed greater order in the light chains of vFP16.02 and vFP20.01 than of vFP1.01 and vFP7.04, especially around the CDR L1 loop. The relative extent of light chain order correlated with the breadth of these antibodies (Fig 4F), suggesting that a more stable light chain interface with Env could lead to increased neutralization breadth.

## Discussion

Overall, these in-depth delineations of the functional epitopes revealed similarities and differences between how VRC34.01 and the vaccine-elicited antibodies recognize Env. Escape from both vFP16.02 and vFP20.01 was mediated predominantly via mutations in the fusion peptide, whereas VRC34.01 neutralization was also affected by mutations at numerous additional sites.

The focused functional interface of the vaccine-elicited antibodies takes on added significance in the context of a unique structural observation: Env-bound vFP16.02 and vFP20.01 have reduced electron density outside of the paratope contacting the fusion peptide’s N-terminus, indicating greater conformational heterogeneity. To further investigate this phenomenon, we generated of two more vFP1 class antibody cryo-EM structures. Intriguingly, these antibodies also displayed internal disorder, and the extent of relative light chain order correlated with breadth (Fig 4F). Increased light chain disorder may indicate weaker interactions with Env gp120. Since our mutational antigenic profiling clearly shows that VRC34.01 but not vFP16.02 or vFP20.01 has functional interactions outside the FP, we hypothesize that additional interactions with other regions of Env may increase the breadth of vFP antibodies. Consistent with this idea, the breadth of vFP antibodies is correlated with their affinity to Env trimer [11]. Excitingly, our functional mapping does show modest but reproducible selection for escape mutations from vFP16.02 and vFP20.01 at site 85 in gp120 (Fig 2), which stacks against their CDR L3 loops (Fig S11). Thus, it does seem possible for the light chain of vaccine-elicited antibodies to derive advantageous interactions from this region of gp120.

Therefore, mapping *all* mutations that affect neutralization by infection and vaccine-elicited antibodies may help identify potentially desirable interactions to target with trimer boosts or rationale immunogen design. However, such interactions could increase affinity at the cost of reduced breadth if the additional Env contacts occur at variable sites. Fortunately, most of the VRC34.01 escape mutations that that are outside of the fusion peptide are at sites largely conserved in nature (Fig S12), suggesting that this tradeoff can be minimized.

An alternative or parallel approach to increasing the breadth of fusion-peptide vaccine responses is to tailor the focused recognition of the fusion peptide to accommodate greater sequence diversity within this region [11]. This goal might be accomplished by immunizing with peptide immunogen pools that incorporate naturally found diversity within these sites [11]. Our data suggest that a peptide immunogen cocktail designed to elicit antibodies similar to vFP16.02 and vFP20.01 would need only to cover diversity at sites 512-516. Further, rather than designing cocktails that cover *all naturally occurring diversity* at sites 512-516, our data allows for the design peptide cocktails that cover only those naturally occurring amino acid polymorphisms *that mediate escape from vFP antibodies*.

In conclusion, our mutational antigenic profiling of vFP antibodies and the template VRC34.01 bnAb, combined with analyses of additional vFP antibody structures, have provided insights that can help refine vaccination regimens. We precisely mapped previously unappreciated interactions between Env and the template bnAb VRC34.01, and our functional and structural analyses suggest that these interactions may help shape the breadth or potency of anti-FP antibodies. Therefore, these complete maps of viral escape provide a detailed atlas to guide subsequent rounds of structure-based vaccine design.

## Materials and Methods

### Antibody production

Antibodies were produced as described in Xu et al (2018) [11].

### Generation of mutant virus libraries

The generation and characterization of the BG505 mutant proviral DNA libraries, and the resulting functional mutant virus libraries have been described in detail previously [14,15]. Briefly, proviral DNA libraries that contained randomized codon-level mutations at sites 31-702 (HXB2 numbering) of BG505.W6M.C2.T332N *env* were independently generated. These, and accompanying wildtype controls, were independently transfected into 293T cells (ATCC). The transfection supernatant was passaged at a low MOI (0.01 TZM-bl infectious units/cell) in SupT1.CCR5 cells (James Hoxie), and the resulting genotype-phenotype linked mutant virus library was concentrated via ultracentrifugation. Env residues are numbered according to HXB2 reference numbering throughout.

### Mutational antigenic profiling

We have previously described Env mutational antigenic profiling using BF520 Env libraries [13]; a similar approach was taken here. Briefly, 5×10^5^ infectious units of two independent mutant virus libraries were neutralized with VRC34.01, vFP16.02, or vFP20.01 at an ~IC_95_ concentration (33 ug/mL, 500 ug/mL, or 500ug/mL of antibody, respectively) for one hour. Neutralized libraries were then infected into 1×10^6^ SupT1.CCR5 cells in R10 (RPMI [GE Healthcare Life Sciences; SH30255.01], supplemented with 10% FBS, 1% 200 mM L-glutamine, and 1% of a solution of 10,000 units/mL penicillin and 10,000 μg/mL streptomycin), in the presence of 100ug/mL DEAE-dextran added. Three hours post infection, cells were spun down and resuspended in 1 mL fresh R10 without DEAE-dextran. At 12 hours post infection, cells were washed with PBS and non-integrated viral cDNA was isolated via miniprep. In parallel to each antibody selection, each mutant virus library was also subjected to a mock selection, and four 10-fold serial dilutions of each mutant virus library were also used to infect 1×10^6^ cells to serve as an infectivity standard curve. Selected and mock-selected viral cDNA was then sequenced with a barcoded subamplicon sequencing approach as previously described [14]. This approach involves introducing unique molecular identifiers during the Illumina library prep in order to further reduce the sequencing error rate.

The relative amount of viral genome that successfully entered cells was measured using *pol* qPCR[16]. A qPCR standard curve was generated based on each mutant virus library’s infectivity standard curve, from which the % neutralization for each antibody selected sample was calculated.

### Analysis of deep sequencing data

We used dms_tools2 version 2.2.4 (https://jbloomlab.github.io/dms_tools2/) to analyze the deep sequencing data and calculate the differential selection [17]. The differential selection statistic has been described in detail [13,18] and is further documented at https://jbloomlab.github.io/dms_tools2/diffsel.html. Sequencing of wildtype proviral DNA plasmid was used as the error control during the calculation of differential selection. Differential selection was visualized on logoplots rendered by dms_tools2 using weblogo [19] and ggseqlogo [20].

### Cryo-EM

HIV-1 Env BG505 DS-SOSIP and antibody Fab fragments were prepared as described previously [21]. For preparing the vFP1.01-BG505 DS-SOSIP-VRC03-PGT122 and the vFP7.04-BG505 DS-SOSIP-VRC03-PGT122 complexes, 4-5-fold molar excess of antibody Fab fragments of vFP1.01 and vFP7.04, respectively, as well as of VRC03 and PGT122, were incubated with HIV-1 Env BG505 DS-SOSIP at a final concentration of 0.3 mg/mL. To prevent aggregation during vitrification, 0.085 mM dodecyl-maltoside (DDM) was added to the sample, followed by vitrification using a semi-automated Spotiton V1.0 robot [22,23]. The grids used were specially designed Nanowire self-blotting grids with a Carbon Lacey supporting substrate [24]. Sample was dispensed onto these nanowire grids using a picoliter piezo dispensing head. A total of ~5 nL sample was dispensed in a stripe across each grid, followed by a pause of a few milliseconds, before the grid was plunged into liquid ethane.

Data was acquired using the Leginon system [25] installed on a Titan Krios electron microscope operating at 300kV and fitted with Gatan K2 Summit direct detection device. The dose was fractionated over 50 raw frames and collected over a 10-s exposure time. Individual frames were aligned and dose-weighted. CTF was estimated using the GCTF package [26]. Particles were picked using DoG Picker [27] within the Appion pipeline [28]. Particles were extracted from the micrographs using RELION [29]. 2D classification, ab initio reconstruction, heterogeneous refinement, and final map refinement were performed using cryoSparc [30]. Postprocessing was performed within RELION.

Fits of the Env trimer and Fab co-ordinates to the cryo-EM reconstructed maps were performed using UCSF Chimera [31]. The coordinates were refined by an iterative process of manual fitting using Coot [32] and real space refinement within Phenix [33]). Molprobity [34] and EMRinger [35] were used to check geometry and evaluate structures at each iteration step. Figures were generated in UCSF Chimera and Pymol (http://www.pymol.org). Electron density quantification and map-fitting cross correlations were calculated within UCSF Chimera. For electron density calculations maps were postprocessed using the larger mask used for refinement in cryoSparc. This mask encompassed the entire complex, including the more disordered antibody constant domains. The maps were then low pass filtered to 4.5 Å in EMAN2 before performing the electron density quantification calculations using the “Values at atom positions” function in Chimera. The electron density at the atom positions for the fitted models were then summed over the heavy and light chains (variable domains only). Map-to-model FSC curves were generated using EMAN2.

### Resistance analysis of 208-pseudovirus panel

We calculated the associations between the sequence variability of FP sequences at sites 512-519 and the 208-pseudovirus panel neutralization data using an in-house version of the approach implemented in R package SeqFeatR^19^. The resulting P-values were corrected using multiple testing method Holm.

### Data availability and source code

Open-source software that recapitulates the mutational antigenic profilng analysis, starting with the deep sequencing data through the calculation of selection metrics, is available at https://jbloomlab.github.io/dms_tools2/. The entire computational analysis, starting with downloading the deep sequencing reads through generating figures, is available as an executable IPython Notebook (File S1) and at https://github.com/jbloomlab/MAP_Vaccine_FP_Abs. Illumina deep sequencing reads were deposited into the NCBI SRA as SRR6429862-SRR6429875.

The cryo-EM maps and fitted coordinates for vFP1.01-BG505 DS-SOSIP-VRC03-PGT122 and vFP7.04-BG505 DS-SOSIP-VRC03-PGT122 have been deposited to the RCSB database with accession numbers EMD-7622/PDB 6CUF and EMD-7621/PDB 6CUE, respectively. Preliminary validation reports are included as File S5 and File S6.

## Acknowledgements

We thank the Fred Hutch Genomics Core for performing the deep sequencing.

---

**Fig S1.**
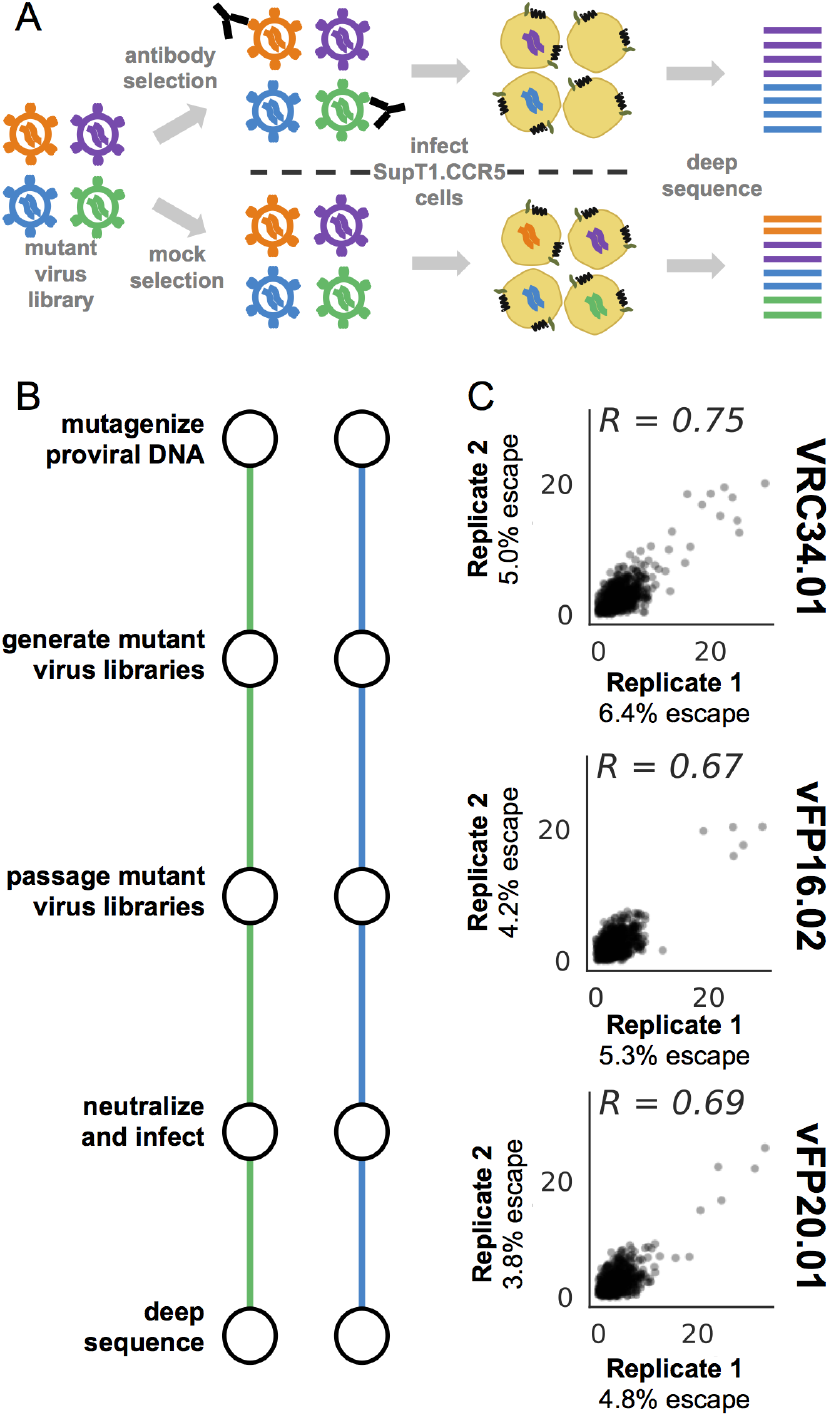
The mutational antigenic profiling pipeline. **A.** Schematic showing the mutational antigenic profiling pipeline. Genotype-phenotype linked mutant virus library, which has undergone functional selection, is neutralized by each antibody before infecting SupT1.CCR5 cells. The mutant virus library is also infected into cells in the absence of antibody selection. Viral cDNA is isolated from each condition and deep sequenced. Antibody escape mutants are enriched in the antibody selected condition relative to the non-selected control. **B.** The entire mutational antigenic profiling pipeline, including generating independent mutant proviral DNA libraries, was performed in duplicate. **C.** The positive site differential selection between biological replicates was well correlated for each antibody. The axes are labeled with the % remaining infectivity relative to the mock selected library for that replicate, measured via qPCR.

**Fig S2.**
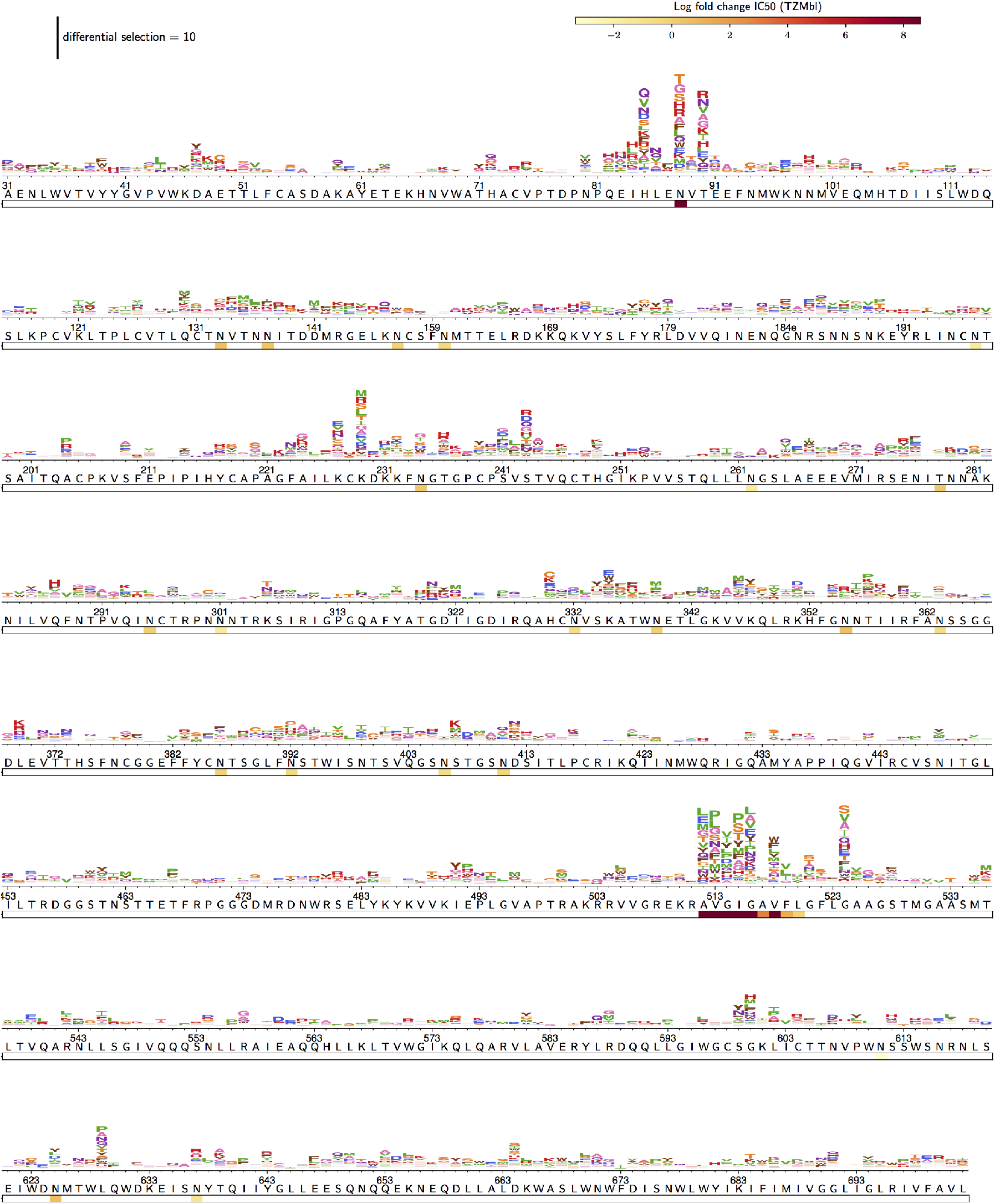
The escape profile for VRC34.01. The height of each amino acid is proportional to the logarithm of the relative enrichment of that mutation in the antibody selected condition relative to the non-selected control. The wildtype BG505.T332N sequence is shown. Underlaid is the original functional mapping data from Kong et al (2016), where a number of mutant BG505 pseudoviruses (lacking the T332N mutation, unlike the BG505 Env used to generate to the mutant libraries) were tested in TZM-bl neutralization assays [5]. If a mutation was tested at a site, then a box is underlaid and colored according to the fold-change in IC_50_ relative to wildtype.

**Fig S3.**
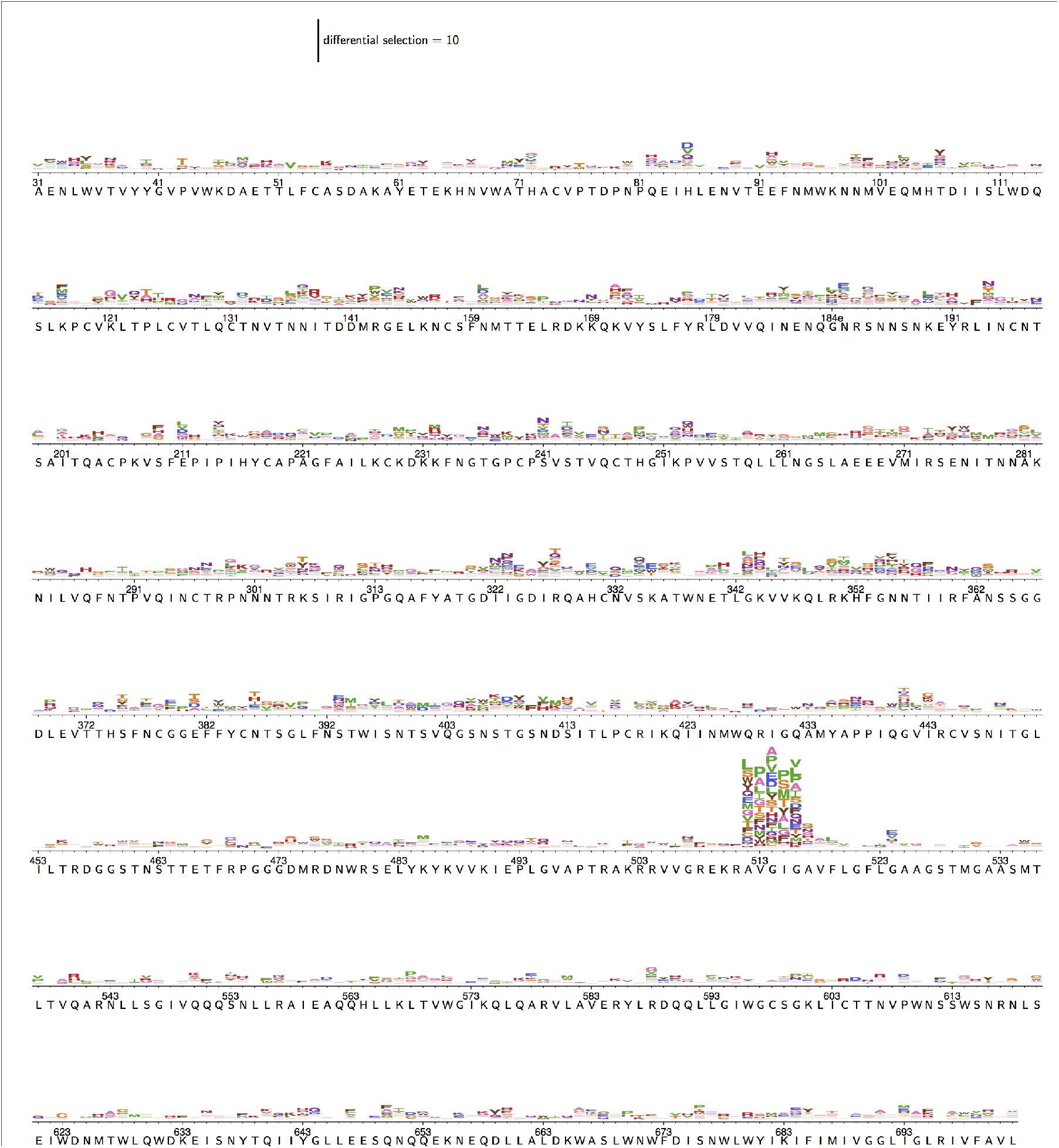
The escape profile for vFP16.02. As described for Fig S2, but for vFP16.02.

**Fig S4.**
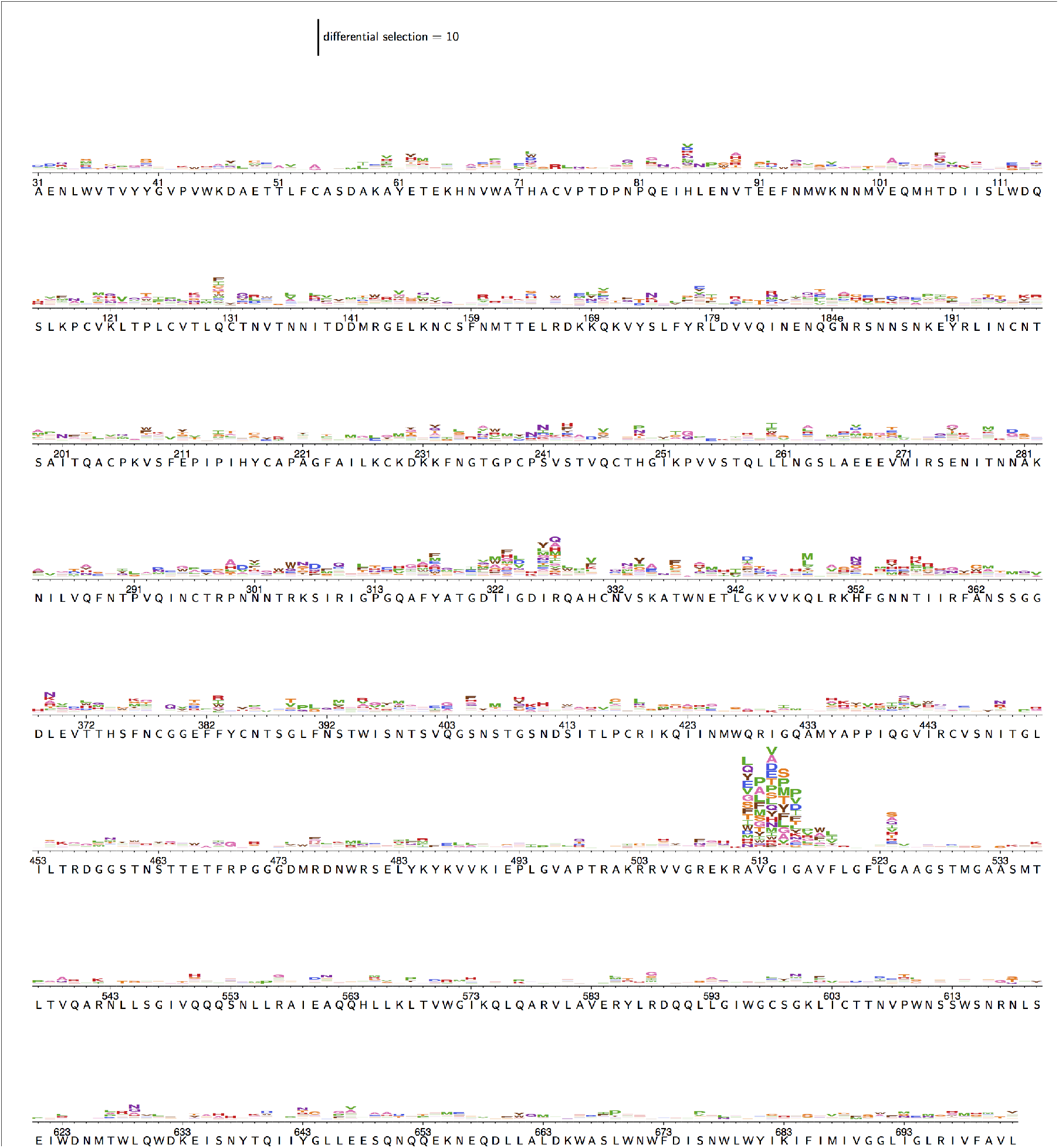
The escape profile for vFP20.01. As described for Fig S2, but for vFP20.01.

**Fig S5.**
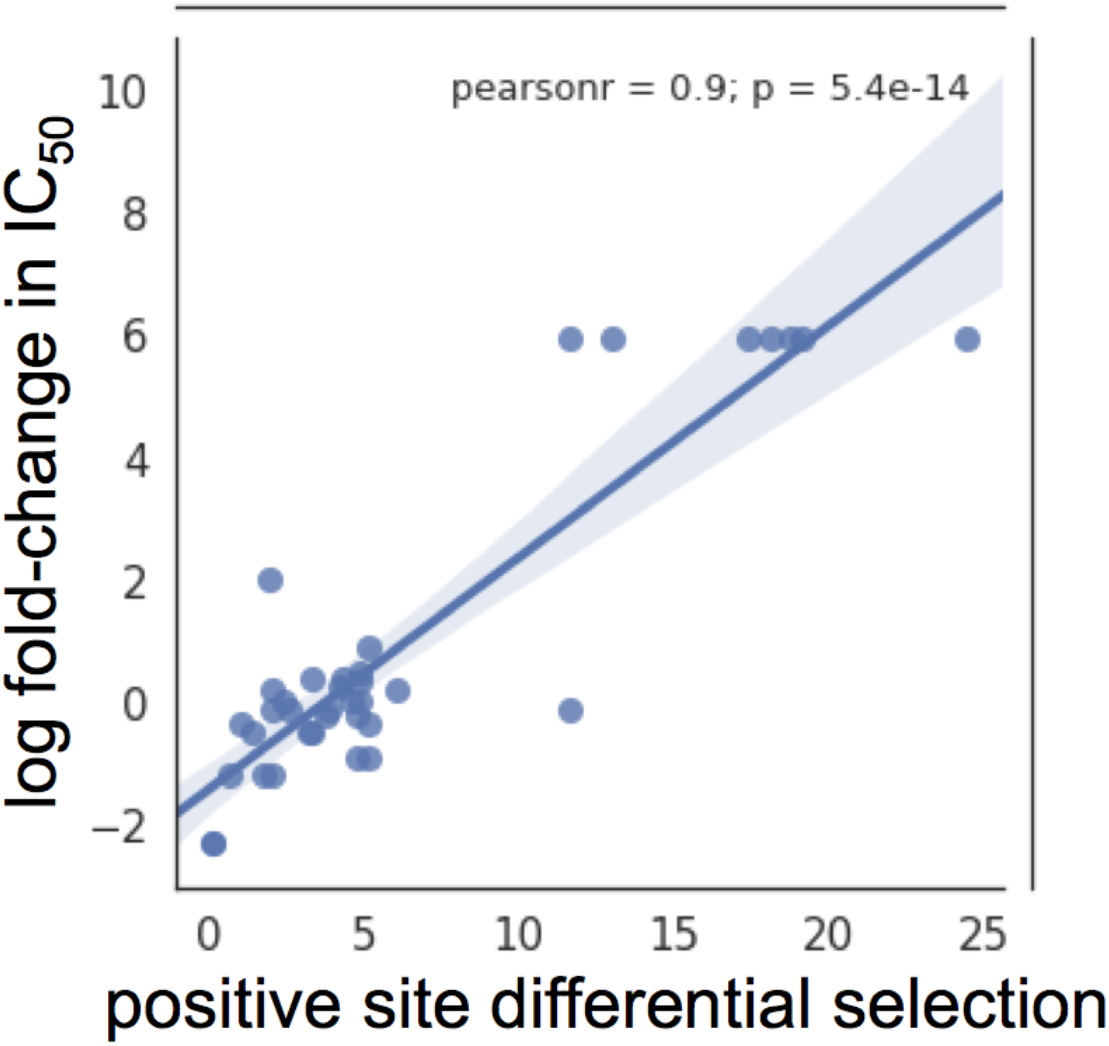
Validation of mutational antigenic profiling. **A.** Correlation between the positive differential selection at a site and the log fold change in IC_50_ from a BG505 pseudovirus bearing a point mutant at that site (TZM-bl data from Kong et al (2016) [5], as described in Fig S2). Note that the TZM-bl data is right censored, and it is possible for a mutation to affect the maximum percent neutralization rather than the IC_50_.

**Tables S1.**
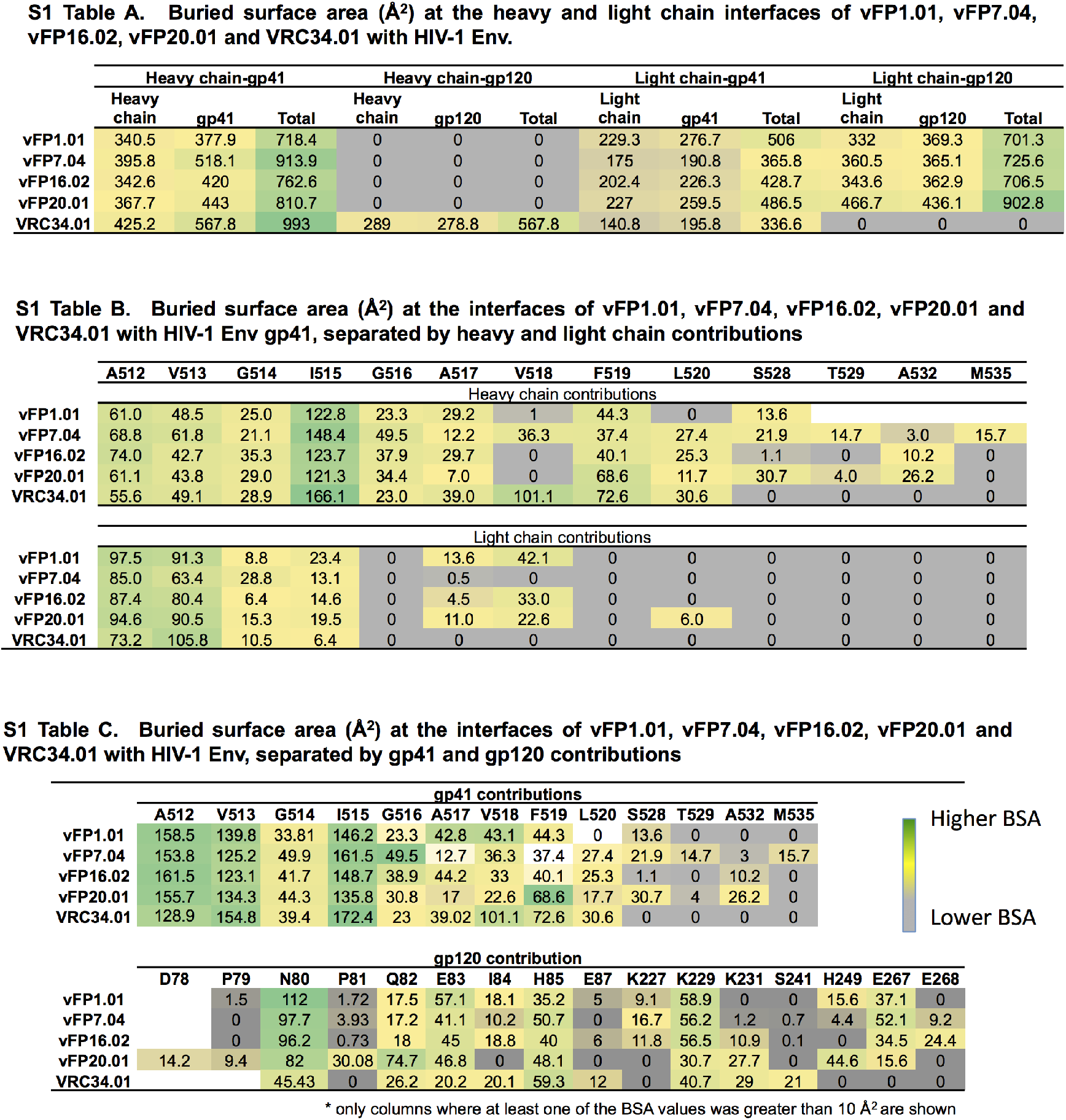
Buried surface area calculations of VRC34.01 and vFP antibodies bound to BG505 Env trimer.

**Table S2.** Cryo-EM data collection, refinement and validation statistics.

|  | vFP1.01-DS-SOSIP-VRC03-PGT122 | vFP7.04-DS-SOSIP-VRC03-PGT122 |
| --- | --- | --- |
|  | EMD-7622; PDB 6CUF | EMD-7621; PDB 6CUE |
| **Data collection and processing** |  |  |
| Magnification | 130000 | 105000 |
| Voltage (kV) | 300 | 300 |
| Electron exposure (e-/Å<sup>2</sup>) | 70.28 | 53.11 |
| Defocus range (μm) | -1.5 - -3.0 | -1.5 - -3.0 |
| Pixel size (Å) | 1.06 | 1.1 |
| Symmetry imposed | C3 | C3 |
| Initial particle images (no.) | 108758 | 152206 |
| Final particle images (no.) | 44652 | 64580 |
| Map resolution (Å) | 4.2 | 4.0 |
| FSC threshold | 0.143 | 0.143 |
| Map resolution range (Å) | 3.9-9.4 | 3.8-8.8 |
| **Refinement** |  |  |
| Initial model used (PDB code) | 6CDI | 6CDI |
| Model resolution (Å) |  |  |
| FSC threshold | 0.5 | 0.5 |
| Model resolution range (Å) | 4.4 | 4.1 |
| Map sharpening *B* factor (Å<sup>2</sup>) | -92.99 | -87.94 |
| Model composition |  |  |
| Non-hydrogen atoms |  |  |
| Protein residues | 3843 | 3849 |
| R.m.s. deviations |  |  |
| Bond lengths (Å) | 0.006 | 0.009 |
| Bond angles (°) | 1.098 | 1.06 |
| Validation |  |  |
| MolProbity score | 1.82 | 1.74 |
| Clashscore | 6.0 | 6.0 |
| Poor rotamers (%) | 0.54 | 0.90 |
| Ramachandran plot |  |  |
| Favored (%) | 93.20 | 93.79 |
| Allowed (%) | 6.72 | 6.13 |
| Disallowed (%) | 0.08 | 0.08 |

**Fig S6.**
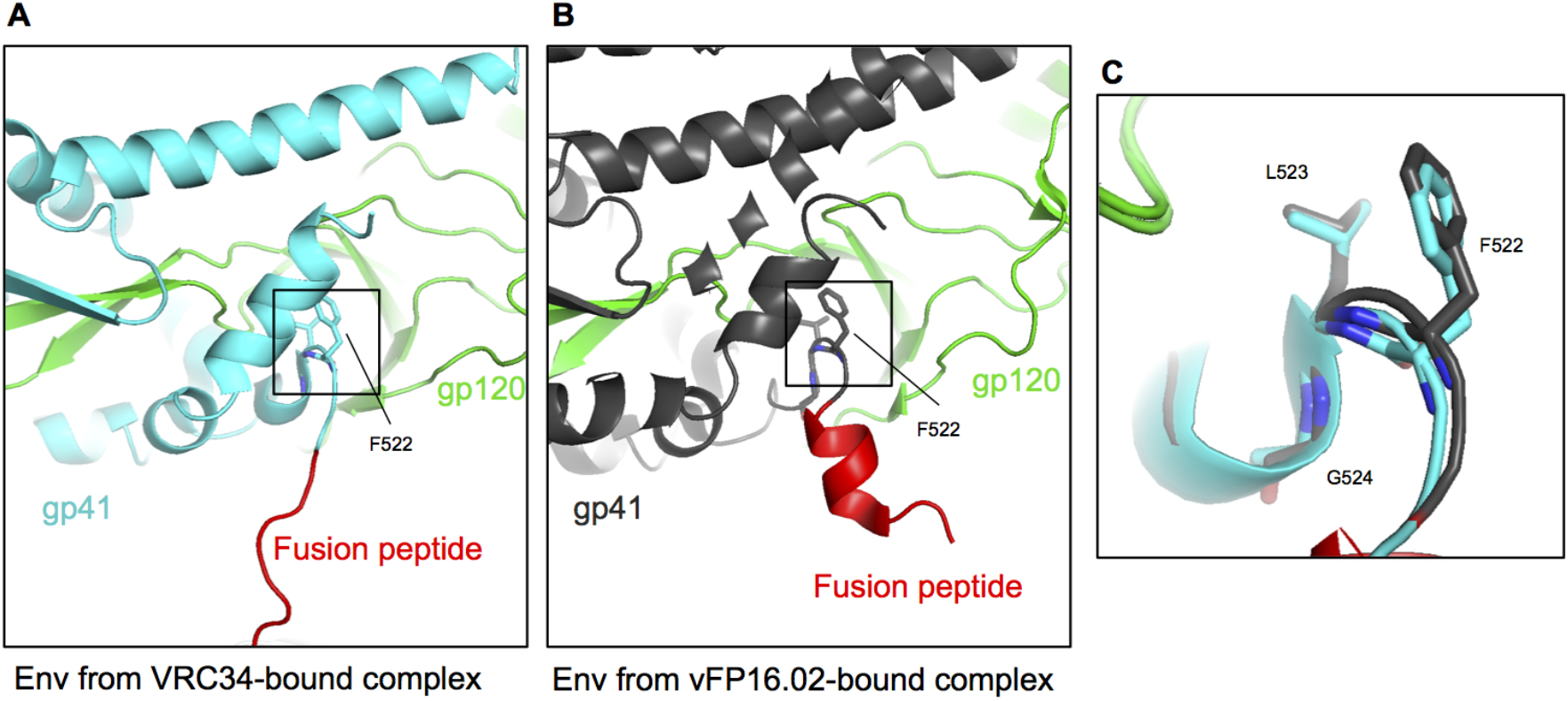
F522 anchors the fusion peptide. A. Env from VRC34-bound complex with gp41 colored cyan, gp120 colored green and the fusion peptide colored red. Residues 522-524 of gp120 are shown in sticks. B. Same as a, but for Env from vFP16.02-bound complex and gp41 shown in dark gray. C. Overlay of VRC34-bound complex and vFP16.02-bound complex.

**Fig S7.**
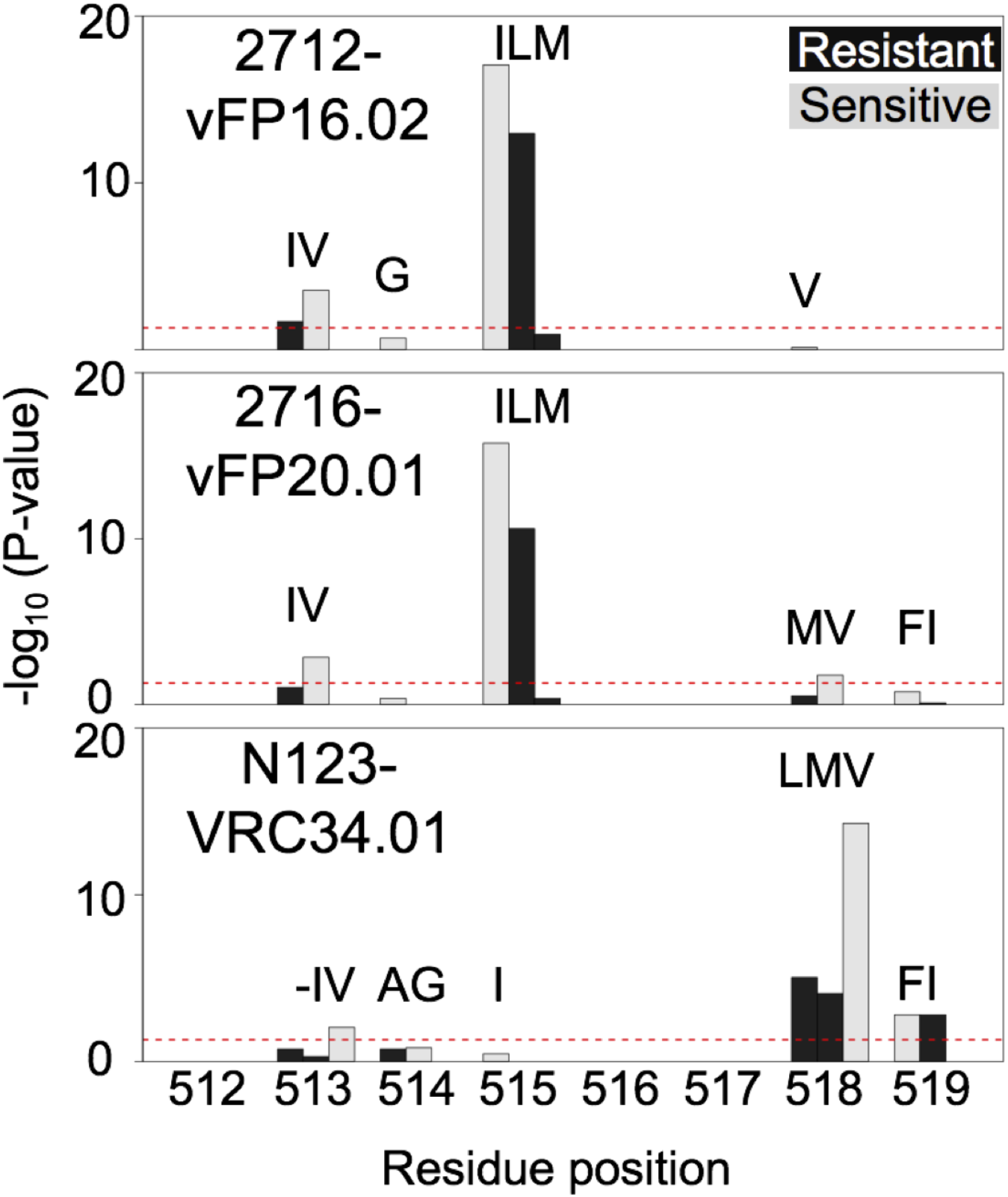
FP sequence association with neutralization for vFP16.02, vFP20.01, and VRC34.01, based on a panel of 208 Env pseudoviruses. Bar plot shows the P-values of the association (calculated by Fisher’s exact test) between amino acid variation and neutralization at residues 512-519. Black bars highlight amino acid and site combinations that significantly associate with resistance to antibodies, while light-gray bars highlight sensitive ones. Amino acids valine and isoleucine at sites 513 and 515, respectively, are significantly associated with sensitivity, while isoleucine at position 513 and leucine at site 515 are significantly associated with resistance to vFP16.02. In the case of vFP20.01, residues valine, isoleucine and valine at sites 513, 515 and 518, respectively, are associated to sensitivity to vFP20.01, with only leucine at position 515 being significantly associated to resistance. For VRC34, valine at positions 513 and 518 as well as phenylalanine at site 519 are significantly associated to sensitivity to VRC34, while leucine and methionine at site 518, and isoleucine at site 519 are associated with resistance. A dotted red line corresponds to the adjusted P-value threshold of 0.05.

**Fig S8.**
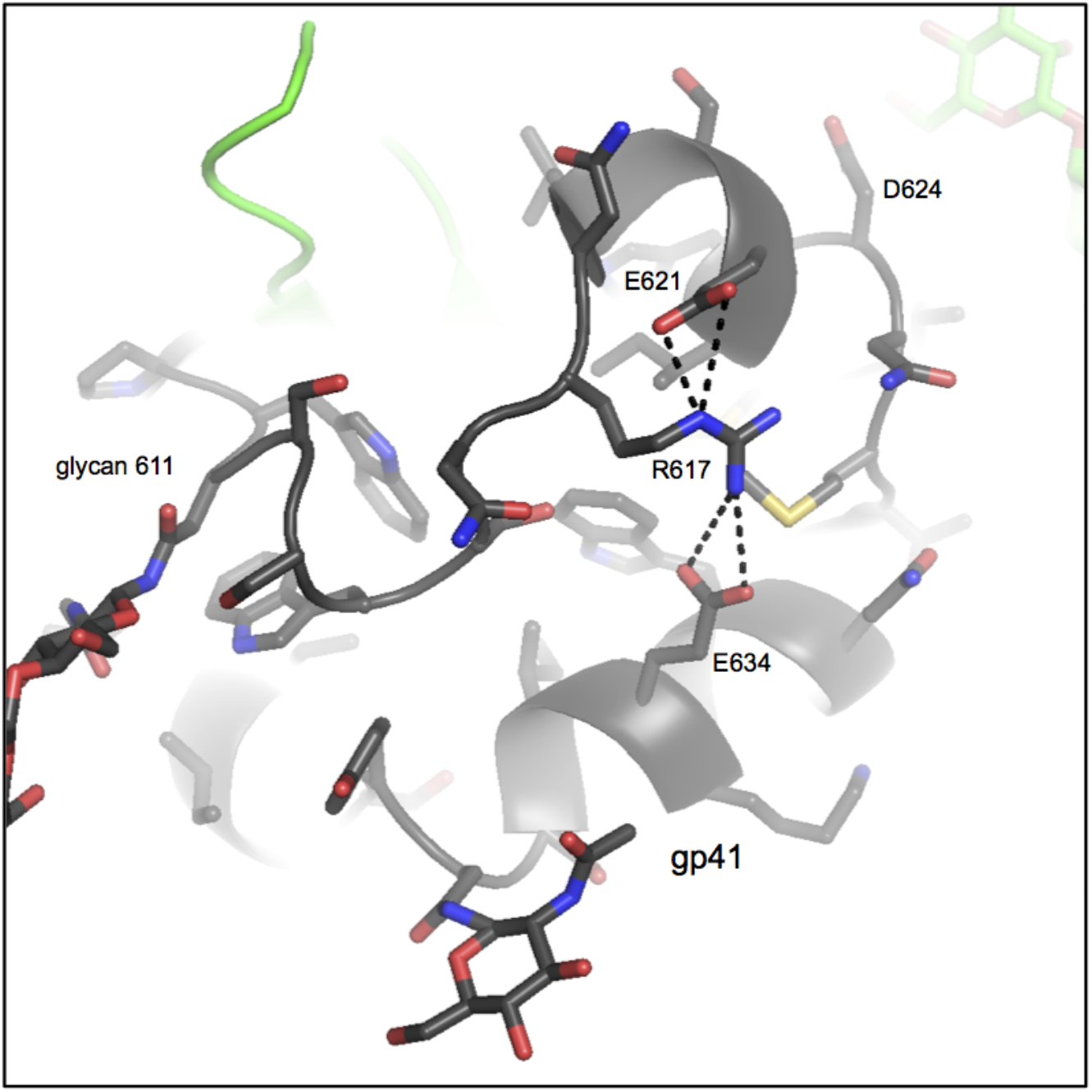
A surface salt bridge network in gp41. Zoom-in of gp41 region showing a surface salt bridge network formed by residues E621, R617 and E634.

**Fig S9.**
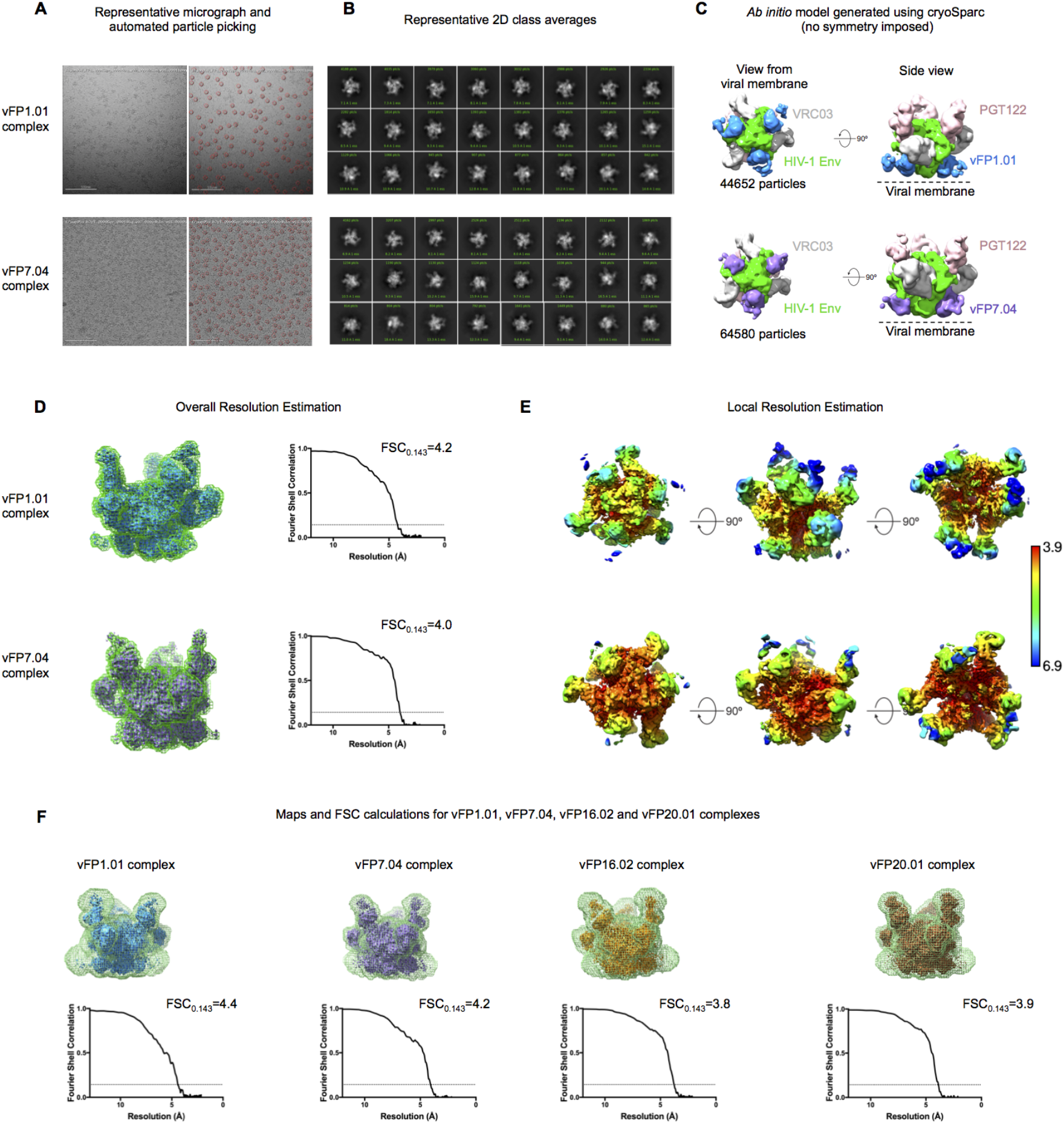
Cryo-EM structural details for vFP1.01-DS-SOSIP-VRC03-PGT122 and vFP7.04-BG505 DS-SOSIP-VRC03-PGT122 complexes. **A.** Representative micrograph (left) with particles picked with DogPicker shown in red circles (right). **B.** Representative 2D class averages. **C.** Ab initio models generated using cryoSparc. **D.** Refined map in blue (for vFP1.01 complex) and purple (for vFP7.04 complex) with the mask used in RELION postprocessing shown in green. FSC plots and resolution values reported by RELION according to the gold standard FSC_0.143_ criterion (FSC_0.143_ shown as dotted line. **E.** Local resolution estimation. **F.** For density quantifications we used maps that were postprocessed using expanded solvent masks that encompassed the entire complex, including the more disordered constant domains. The FSC plots and overall resolutions of these maps are shown. The maps were then low-pass filtered to 4.5 Å for the density quantifications.

**Fig S10.**
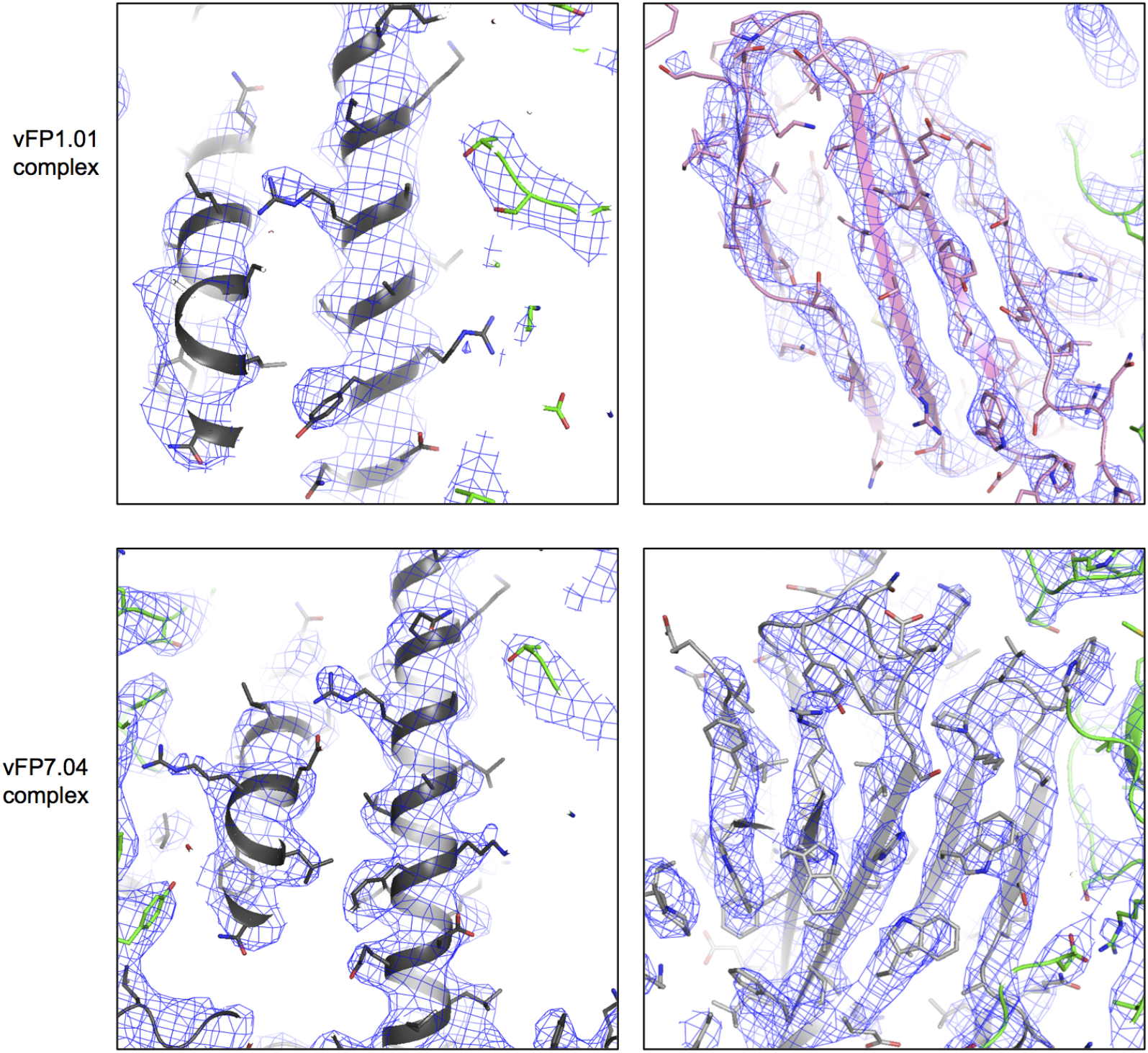
Representative electron density snapshots for the vFP1.01-DS-SOSIPVRC03-PGT122 and vFP7.04-BG505 DS-SOSIP-VRC03-PGT122 complexes. Electron density is shown as a blue mesh and the fitted model is shown as a cartoon with residue side-chains shown as sticks.

**Fig S11.**
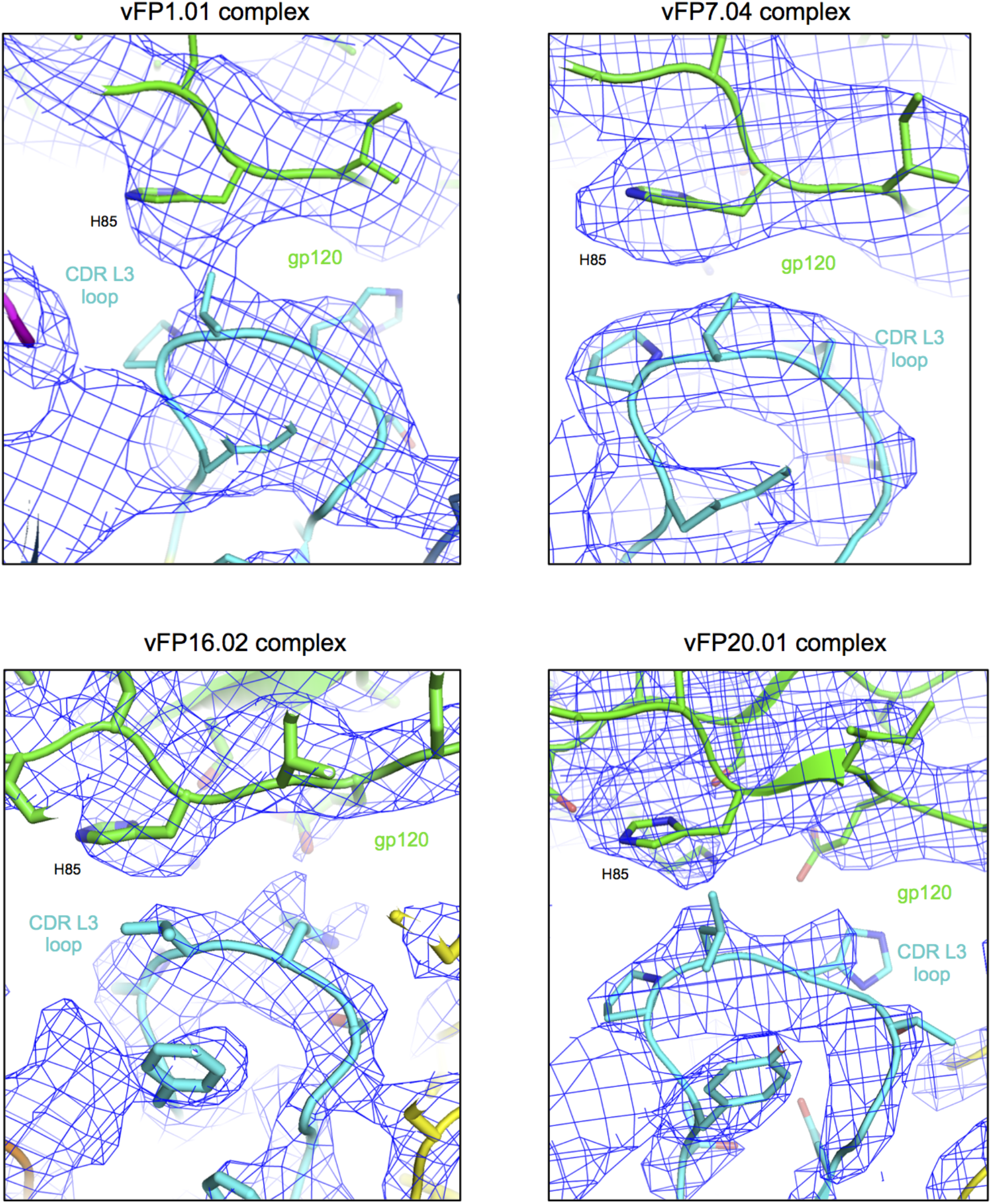
Interaction of His 85 with CDR L3 loop of vFP Mabs.

**Fig S12.**
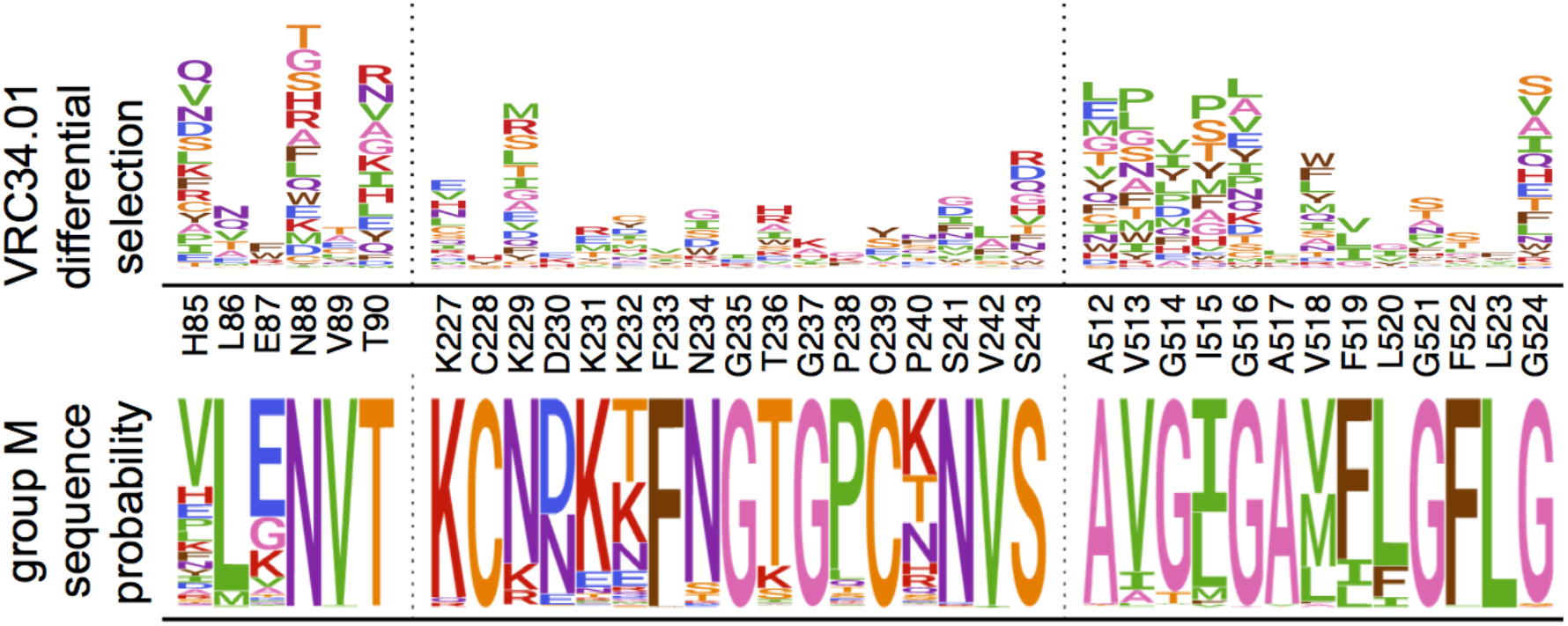
Sequence conservation of the VRC34.01 functional epitope in nature. The VRC34.01 escape profile is shown as in Fig 2A. Beneath, the sequence conservation among group M sequence LANL Web Alignment is shown.

## Supplemental Files

**File S1 | Mutational Antigenic Profiling IPython notebook analysis** This zipped file contains the entire mutational antigenic profiling analysis. There is an html document of the completed analysis, an executable IPython notebook, and all of the necessary input data to execute the analysis.

**File S2 | VRC34.01 differential selection** The VRC34.01 differential selection values for all measured mutations. The format is described at https://jbloomlab.github.io/dms_tools2/diffsel.html and in Dingens et al 2017 [13].

**File S3 | vFP16.02 differential selection** The vFP16.02 differential selection values for all measured mutations.

**File S4 | vFP20.01 differential selection** The vFP20.01 differential selection values for all measured mutations.

**File S5 | wwPDB/EMDataBank EM Map/Model Validation Report for vFP1.01-BG505 DS-SOSIP-VRC03-PGT122** Preliminary validation report for the cryo-EM maps and fitted coordinates for vFP1.01-BG505 DS-SOSIP-VRC03-PGT122.

**File S6 | wwPDB/EMDataBank EM Map/Model Validation Report for vFP7.04-BG505 DS-SOSIP-VRC03-PGT122** Preliminary validation report for the cryo-EM maps and fitted coordinates for vFP7.04-BG505 DS-SOSIP-VRC03-PGT122.

## References

1. Burton DR, Ahmed R, Barouch DH, Butera ST, Crotty S, Godzik A, et al. A Blueprint for HIV Vaccine Discovery. Cell Host Microbe. 2012;12: 396–407. doi:10.1016/j.chom.2012.09.008

2. Kwong PD, Mascola JR. Human antibodies that neutralize HIV-1: identification, structures, and B cell ontogenies. Immunity. 2012;37: 412–25. doi:10.1016/j.immuni.2012.08.012

3. Wu X, Kong X-P. Antigenic landscape of the HIV-1 envelope and new immunological concepts defined by HIV-1 broadly neutralizing antibodies. Curr Opin Immunol. 2016;42: 56–64. doi:10.1016/j.coi.2016.05.013

4. McCoy LE, Burton DR. Identification and specificity of broadly neutralizing antibodies against HIV. Immunol Rev. 2017;275: 11–20. doi:10.1111/imr.12484

5. Kong R, Xu K, Zhou T, Acharya P, Lemmin T, Liu K, et al. Fusion peptide of HIV-1 as a site of vulnerability to neutralizing antibody. Science (80-). 2016;352: 423–426. doi:10.1126/science.aae0474

6. Wibmer CK, Gorman J, Ozorowski G, Bhiman JN, Sheward DJ, Elliott DH, et al. Structure and Recognition of a Novel HIV-1 gp120-gp41 Interface Antibody that Caused MPER Exposure through Viral Escape. PLoS Pathog. 2017;13: e1006074. doi:10.1371/journal.ppat.1006074

7. Van Gils MJ, Van Den Kerkhof TLGM, Ozorowski G, Cottrell C a, Sok D, Pauthner M, et al. An HIV-1 antibody from an elite neutralizer implicates the fusion peptide as a site of vulnerability. Nat Microbiol. 2016; 1–10. doi:10.1038/NMICROBIOL.2016.199

8. Blattner C, Lee JH, Sliepen K, Derking R, Falkowska E, de la Peña AT, et al. Structural Delineation of a Quaternary, Cleavage-Dependent Epitope at the gp41-gp120 Interface on Intact HIV-1 Env Trimers. Immunity. 2014;40: 669–680. doi:10.1016/j.immuni.2014.04.008

9. Falkowska E, Le KM, Ramos A, Doores KJ, Lee JH, Blattner C, et al. Broadly Neutralizing HIV Antibodies Define a Glycan-Dependent Epitope on the Prefusion Conformation of gp41 on Cleaved Envelope Trimers. Immunity. 2014;40: 657–668. doi:10.1016/j.immuni.2014.04.009

10. Friedman J, Alam SM, Shen X, Xia SM, Stewart S, Anasti K, et al. Isolation of HIV-1-neutralizing mucosal monoclonal antibodies from human colostrum. PLoS One. 2012;7. doi:10.1371/journal.pone.0037648

11. Kai Xu, Acharya P, Kong R, Cheng C, Chuang G-Y, Liu K, et al. Epitope-based vaccine design yields fusion peptide-directed antibodies that neutralize diverse strains of HIV-1. bioRxiv. 2018. doi: 10.1101/306282

12. Trkola a, Purtscher M, Muster T, Ballaun C, Buchacher a, Sullivan N, et al. Human monoclonal antibody 2G12 defines a distinctive neutralization epitope on the gp120 glycoprotein of human immunodeficiency virus type 1. J Virol. 1996;70: 1100–1108.

13. Dingens AS, Haddox HK, Overbaugh J, Bloom JD. Comprehensive Mapping of HIV-1 Escape from a Broadly Neutralizing Antibody. Cell Host Microbe. 2017;21: 1–11. doi:10.1016/j.chom.2017.05.003

14. Haddox HK, Dingens AS, Hilton SK, Overbaugh J, Bloom JD. Mapping mutational effects along the evolutionary landscape of HIV envelope. Elife. 2018;7. doi:10.7554/eLife.34420

15. Haddox HK, Dingens AS, Bloom JD. Experimental Estimation of the Effects of All Amino-Acid Mutations to HIV’s Envelope Protein on Viral Replication in Cell Culture. PLOS Pathog. 2016;12: e1006114. doi:10.1371/journal.ppat.1006114

16. Benki S, McClelland RS, Emery S, Baeten JM, Richardson BA, Lavreys L, et al. Quantification of genital human immunodeficiency virus type 1 (HIV-1) DNA in specimens from women with low plasma HIV-1 RNA levels typical of HIV-1 nontransmitters. J Clin Microbiol. 2006;44: 4357–4362. doi:10.1128/JCM.01481-06

17. Bloom JD. Software for the analysis and visualization of deep mutational scanning data. BMC Bioinformatics. 2015;16: 1–13. doi:10.1186/s12859-015-0590-4

18. Doud MB, Hensley SE, Bloom JD. Complete mapping of viral escape from neutralizing antibodies. PLOS Pathog. 2017;13: e1006271. doi:10.1371/journal.ppat.1006271

19. Crooks GE, Hon G, Chandonia J-M, Brenner SE. WebLogo: a sequence logo generator. Genome Res. 2004;14: 1188–90. doi:10.1101/gr.849004

20. Wagih O. Ggseqlogo: A versatile R package for drawing sequence logos. Bioinformatics. 2017;33: 3645–3647. doi:10.1093/bioinformatics/btx469

21. Kwon Y Do, Pancera M, Acharya P, Georgiev IS, Crooks ET, Gorman J, et al. Crystal structure, conformational fixation and entry-related interactions of mature ligand-free HIV-1 Env. Nat Struct Mol Biol. 2015;22: 522–531. doi:10.1038/nsmb.3051

22. Razinkov I, Dandey VP, Wei H, Zhang Z, Melnekoff D, Rice WJ, et al. A new method for vitrifying samples for cryoEM. J Struct Biol. 2016;195: 190–198. doi:10.1016/j.jsb.2016.06.001

23. Dandey VP, Wei H, Zhang Z, Tan YZ, Acharya P, Eng ET, et al. Spotiton: New features and applications. J Struct Biol. 2018;202: 161—169. doi:10.1016/j.jsb.2018.01.002

24. Wei H, Dandey VP, Zhang Z, Raczkowski A, Rice WJ, Carragher B, et al. Optimizing “self-wicking” nanowire grids. Journal of Structural Biology. 2018. doi:10.1016/j.jsb.2018.01.001

25. Suloway C, Pulokas J, Fellmann D, Cheng A, Guerra F, Quispe J, et al. Automated molecular microscopy: The new Leginon system. J Struct Biol. 2005;151: 41–60. doi:10.1016/j.jsb.2005.03.010

26. Zhang K. Gctf: Real-time CTF determination and correction. J Struct Biol. 2016;193: 1–12. doi:10.1016/j.jsb.2015.11.003

27. Voss NR, Yoshioka CK, Radermacher M, Potter CS, Carragher B. DoG Picker and TiltPicker: Software tools to facilitate particle selection in single particle electron microscopy. J Struct Biol. 2009;166: 205–213. doi:10.1016/j.jsb.2009.01.004

28. Lander GC, Stagg SM, Voss NR, Cheng A, Fellmann D, Pulokas J, et al. Appion: An integrated, database-driven pipeline to facilitate EM image processing. J Struct Biol. 2009;166: 95–102. doi:10.1016/j.jsb.2009.01.002

29. Scheres SHW. RELION: Implementation of a Bayesian approach to cryo-EM structure determination. J Struct Biol. 2012;180: 519–530. doi:10.1016/j.jsb.2012.09.006

30. Punjani A, Rubinstein JL, Fleet DJ, Brubaker MA. CryoSPARC: Algorithms for rapid unsupervised cryo-EM structure determination. Nat Methods. 2017;14: 290–296. doi:10.1038/nmeth.4169

31. Pettersen EF, Goddard TD, Huang CC, Couch GS, Greenblatt DM, Meng EC, et al. UCSF Chimera—A Visualization System for Exploratory Research and Analysis. J Comput Chem. 2004;25: 1605–1612. doi:10.1002/jcc.20084

32. Emsley P, Cowtan K. Coot: Model-building tools for molecular graphics. Acta Crystallogr Sect D Biol Crystallogr. 2004;60: 2126–2132. doi:10.1107/S0907444904019158

33. Adams PD, Gopal K, Grosse-Kunstleve RW, Hung LW, Ioerger TR, McCoy AJ, et al. Recent developments in the PHENIX software for automated crystallographic structure determination. Journal of Synchrotron Radiation. 2004. pp. 53–55. doi:10.1107/S0909049503024130

34. Davis IW, Murray LW, Richardson JS, Richardson DC. MolProbity: Structure validation and all-atom contact analysis for nucleic acids and their complexes. Nucleic Acids Res. 2004;32. doi:10.1093/nar/gkh398

35. Barad BA, Echols N, Wang RYR, Cheng Y, Dimaio F, Adams PD, et al. EMRinger: Side chain-directed model and map validation for 3D cryo-electron microscopy. Nat Methods. 2015;12: 943–946. doi:10.1038/nmeth.3541

